# Combined SNP parental haplotyping and intensity analysis identifies meiotic and mitotic aneuploidies and frequent segmental aneuploidies in preimplantation human embryos

**DOI:** 10.1101/2024.11.17.623999

**Authors:** Alan H Handyside, Louise Newnham, Matthew Newnham, Dominika Henning, Jan Velebny, Jan Pozdena, Jindriska Krmelova, Jakub Horak

## Abstract

Genome-wide single nucleotide polymorphism (SNP) genotyping using microarrays and karyomapping (parental haplotyping) is a universal linkage-based method for preimplantation genetic testing of monogenic disease (PGT-M) and identification of chromosome aneuploidies, including meiotic trisomies, monosomies and deletions. Following IVF, embryos are biopsied at the blastocyst stage and several trophectoderm cells removed. Both parents, a close relative of known disease status and the biopsy samples are genotyped and parental haplotypes analysed. Here we extended the method by combining parental haplotyping with SNP intensity analysis. This enables identification of meiotic and mitotic, whole and segmental aneuploidies at high resolution. In 342 cycles of PGT-M in couples with a mean maternal age of 32.9±4.2 (SD), 37% (471/1270) of the biopsy samples were identified as aneuploid with an almost equal number of meiotic and mitotic aneuploidies. Meiotic aneuploidies were predominantly whole chromosome aneuploidies of maternal origin and increased with maternal age. Mitotic aneuploidies (with normal parental haplotype patterns) were mainly segmental imbalances. For PGT of aneuploidies (PGT-A) in infertile couples, identifying meiotic aneuploidies, which are almost all non-viable, provides a valuable option to avoid the discard of embryos with only mitotic aneuploidies of unknown clinical outcome.

## Introduction

Chromosome aneuploidy (abnormal chromosome number) is a major cause of pregnancy failure, miscarriage and rarely, abnormal pregnancy and live births, after normal or assisted conception (Levy *et al*., 2014; Segawa *et al*., 2017). Whilst most aneuploidies are incompatible with live birth, aneuploidies of the small acrocentric chromosomes and the sex chromosomes, are compatible with development to term, though the incidence at birth is rare (0.3-0.5%) (Moorthie *et al*., 2018). Hence, preimplantation genetic testing for aneuploidy (PGT-A) is now widely used to select viable euploid embryos for transfer following IVF and biopsy of a small number of the outer, extraembryonic trophectoderm cells at the blastocyst stage.

Chromosome gains and losses arising from errors during meiosis, prior to fertilisation, result in trisomies and monosomies, respectively, affecting all the cells of the embryo. Most meiotic aneuploidies arise in female meiosis and increase exponentially with advanced maternal age after 35 years of age (Hassold and Hunt, 2001; Nagaoka *et al*., 2012). Segregation errors and other abnormalities of mitosis following fertilisation, however, are also common. Depending on when they occur, mitotic aneuploidies only affect a variable proportion of cells in the embryo leading to chromosome mosaicism. In addition, chromosome breaks, and other abnormal events can cause gain or loss of whole or part of a chromosome arm causing segmental aneuploidies (Vanneste *et al*., 2009; Chavli *et al*., 2024). Depending on the chromosomes involved and the proportion of affected cells, these aneuploidies may contribute to developmental arrest as the embryo makes the transition to zygotic gene expression before the blastocyst stage, which occurs in about half of human embryos following IVF (McCoy *et al*., 2023).

Genome-wide single nucleotide polymorphism (SNP) genotyping using microarrays and karyomapping (parental haplotyping) is a universal linkage-based method for preimplantation genetic testing of monogenic disease (PGT-M) and identification of meiotic trisomies, monosomies and deletions (Handyside *et al*., 2010; Natesan *et al*., 2014). By genotyping both parents and a close relative, typically a child or grandparent, biallelic SNPs are phased at successive loci across each chromosome and haplotypes for each of the four parental chromosomes established. This, together with knowledge of the disease status of the parents and reference, then allows linkage-based testing by comparing the genotype of biopsied cells from each embryo with the reference, in the region of the gene. Furthermore, genome-wide analysis of parental haplotypes allows detection of aneuploidies by the presence of both haplotypes (dual haplotypes) or neither haplotype from one parent, for either the whole chromosome or a segment.

From 2014-2022, a large reference lab completed over 14,600 PGT-M cycles by karyomapping in 8,400 cases, for 4000 mutations and 900 disorders (Schadwell *et al*., 2022). However, chromosome copy number analysis by low read depth next generation sequencing (NGS) was performed in parallel for the identification of aneuploidy in any unaffected embryos.

Microarray-based genome-wide SNP analysis was developed in the early 2000s and is routinely used for high-throughput genome-wide association studies (GWAS) and high-resolution molecular cytogenetics (Mei *et al*., 2000; Rauch *et al*., 2004; Wollstein *et al*., 2007). For cytogenetics, intensity ratio (log R ratio) and B-allele frequency (BAF) plots are used to analyse copy number variation (CNV), loss of heterozygosity (LOH) and mosaicism. If required, parental genotyping then allows the parental origin of any CNVs to be investigated. Several methods have been developed to combine analysis of SNP genotype and intensity data from microarrays for PGT including trisomies of mitotic origin which cannot be detected by genotype analysis alone (Johnson *et al*., 2010; Zamani Esteki *et al*., 2015). However, published clinical data is limited and combining karyomapping with intensity analysis using BAF and log R ratio has only recently been reported (Kubicek *et al*., 2019).

Here we report an analytical method for comprehensive preimplantation genetic testing of monogenic disease, structural chromosome rearrangements and aneuploidy (PGT-M/SR/A) by combining microarray-based SNP genotyping and karyomapping with parental intensity ratio analysis in a single assay. Using this approach for PGT-A does not require a reference to phase the SNP calls and enables the identification of whole chromosome and segmental aneuploidies of both meiotic and mitotic origin at high resolution, together with karyotype-wide abnormalities including those arising from abnormal fertilisation or contamination.

## Results

Over 12 months from February 2023, 342 cycles of IVF and preimplantation genetic testing for monogenic disease and/or structural chromosome rearrangements (PGT-M/SR) by SNP genotyping and karyomapping were carried out in couples with a mean maternal age of 32.9±4.2 (SD) (range 24-43 years). In total, 1409 blastocysts (4.1 per cycle) were biopsied on days 5-7 post insemination and 3-10 trophectoderm cell samples processed for whole genome amplification (WGA) and SNP genotyping.

Following conventional karyomap analysis for the monogenic disease and/or any chromosomal imbalance related to structural rearrangements in the parents (data not shown), combined parental haplotyping (karyomapping) and intensity ratio profiles were analysed in 1279 samples (90.8%) allowing the relative intensity of the paternal and maternal chromosomes to be compared to the corresponding parental haplotype patterns (Figures 1-3). In 130 samples (9.2%), the parental intensities were too variable to provide a reliable comparison (genotyping and karyomapping was still possible in many cases), maternal contamination was detected or WGA failed.

**Figure 1.**
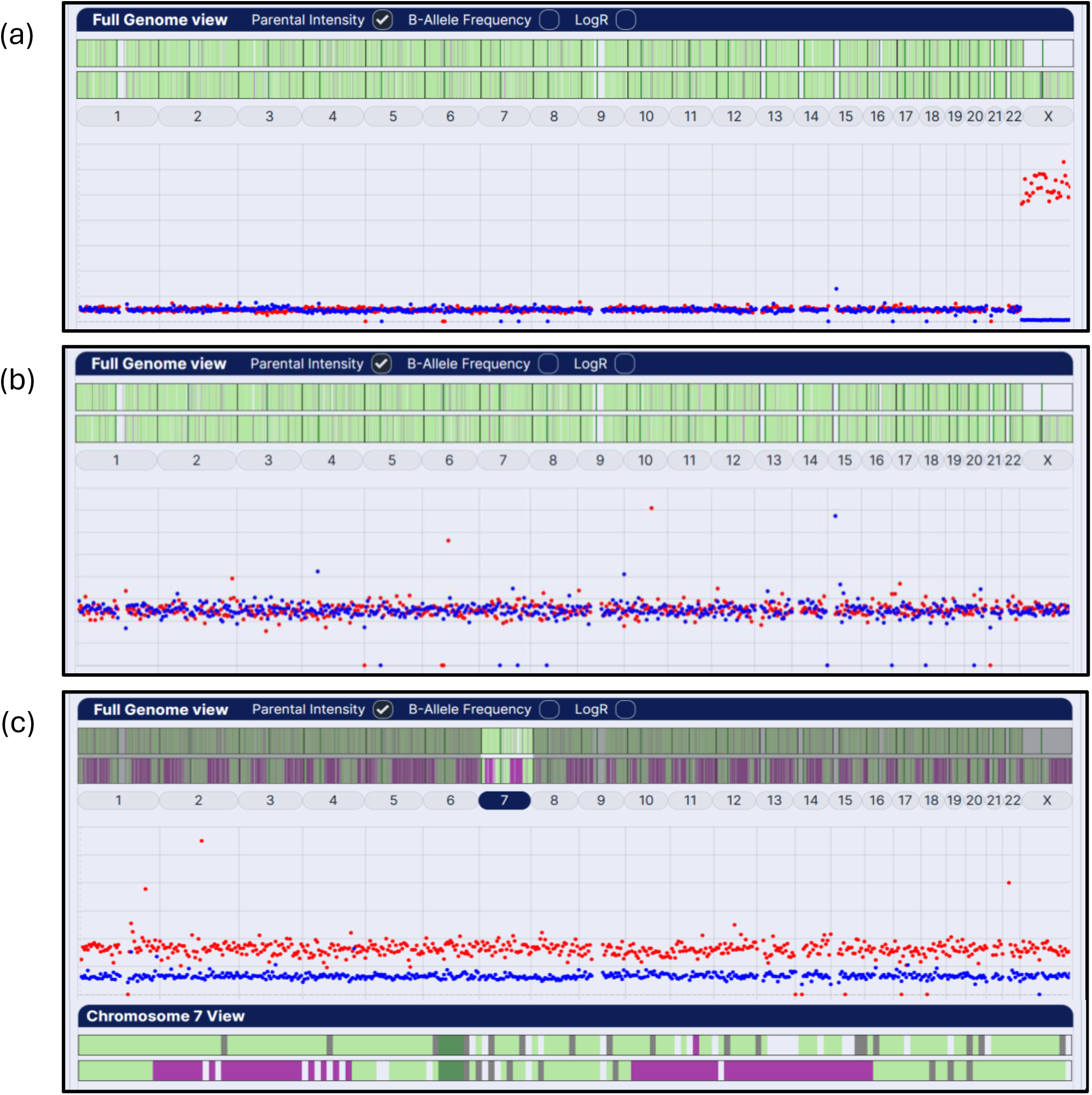
Genome-wide combined parental haplotype and intensity ratio analysis in normal euploid and abnormally fertilised trophectoderm biopsy samples. (a) Normal euploid male (note the closely similar paternal and maternal ratios for chromosomes 1-22 and the low intensity of the paternal X chromosome and the correspondingly elevated maternal intensity ratio), (b) normal euploid female (note the similar ratios of all chromosomes including the paternal and maternal X chromosomes), and (c) a digynic triploid sample from a presumed abnormally fertilised embryo, with elevated maternal intensity ratios and a pattern of dual maternal haplotypes for all chromosomes characteristic of complete failure of the second meiotic division (MII type) as in the example of chromosome 7 with a single maternal haplotype in the pericentromeric region and regions of dual haplotypes on both p and q arms. Genome-wide paternal (top) and maternal (below) haplotypes are displayed across the top of each panel: normal single haplotype (pale green), dual haplotypes (purple) or none (grey) and insufficient SNP data (blank) analysed in successive 1Mb regions across each chromosome with centromeric regions marked in dark green. Paternal (blue) and maternal (red) intensity ratios averaged over successive 5Mb regions of each chromosome are displayed in the scatter plots below.

Excluding nine biopsy samples, 0.7% (9/1279), from abnormally fertilised embryos with karyotype-wide meiotic trisomies or monosomies (six digynic triploids, two diandric triploids and one maternal haploid), combined parental haplotyping and intensity ratio analysis identified 63% (799/1270) of trophectoderm biopsy samples as euploid (no abnormalities detected) and 37% (471/1270) as aneuploid with a total of 664 whole and segmental chromosome aneuploidies, 366 (55%) of meiotic and 298 (45%) of presumed mitotic origin (Table 1; Figures 1 and 2). Maternal meiotic aneuploidies were the most frequent type of abnormality identified and increased significantly in women of advanced maternal age: 11% (96/856) vs 46% (189/414) in ages ≤34 and ≥35 years, respectively (p <0.0001). Maternal meiotic whole chromosome aneuploidies were more frequent in the smaller mainly acrocentric chromosomes, with chromosomes 16 and 22 being the most frequently affected (Figure 3a). Segregation errors in the first meiotic division characterised by dual haplotypes in the pericentromeric and distal regions of affected chromosomes (MI type) were more frequent than those with dual haplotype regions restricted to the chromosome arms (MII type), 57% (73/128), and this trend was more pronounced in the smaller chromosomes (Figure 3b). In comparison, paternal meiotic aneuploidies were relatively rare, most were monosomies, and only four paternal trisomies were identified.

**Figure 2.**
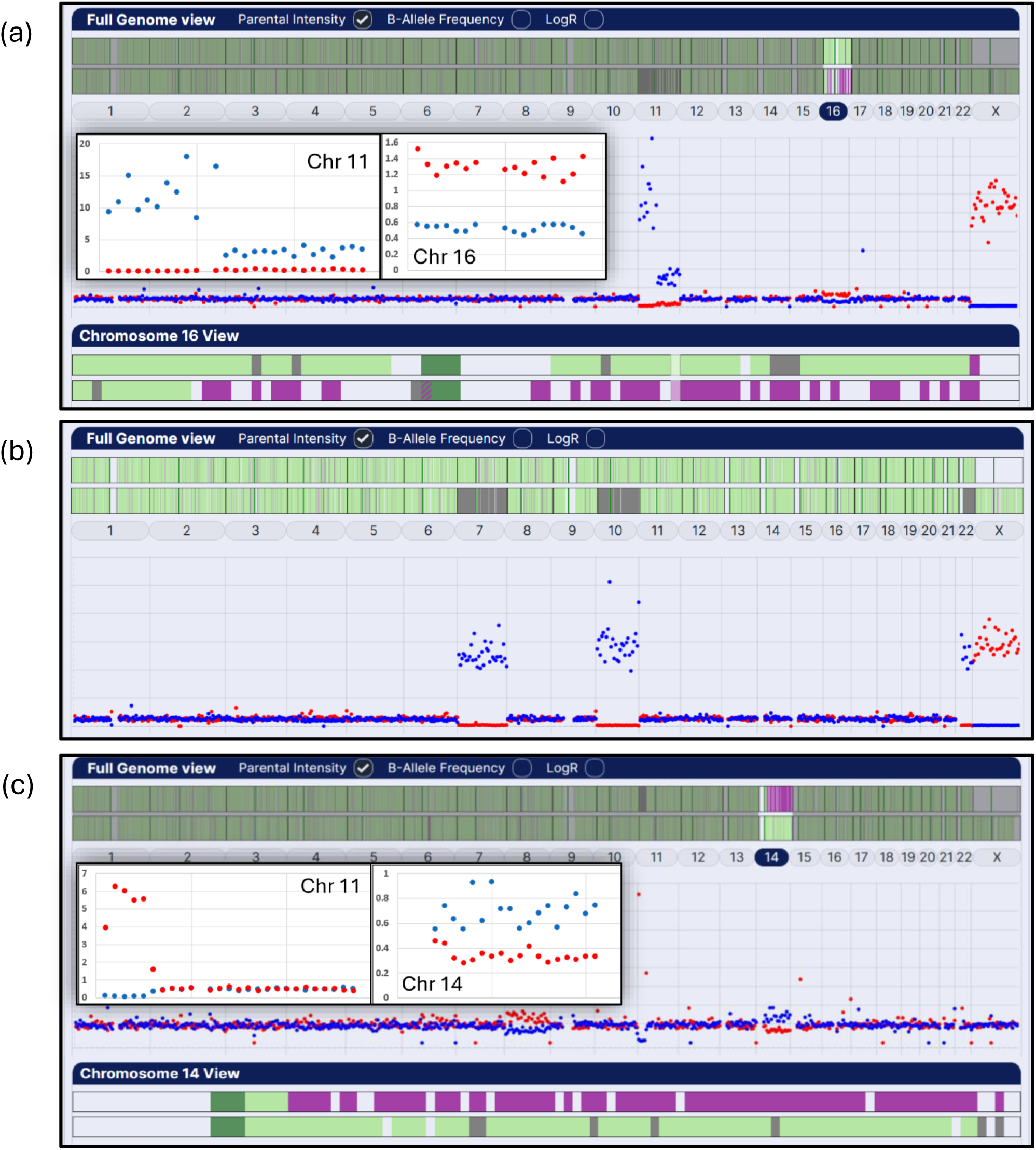
Genome-wide combined parental haplotype and intensity ratio analysis in trophectoderm biopsy samples with various types of aneuploidies. (a) aneuploid sample with a complex segmental aneuploidy of chromosome 11 (loss of maternal p arm with mitotic gain of whole paternal chromosome and maternal trisomy 16 with dual haplotypes across the pericentromeric region typical of a segregation error in the first meiotic division (MI type), (b) maternal monosomies of chromosomes 7, 10 and 22, and (c) paternal loss of chromosome 11 pter and gain of chromosome 14 qter and a mitotic gain of maternal chromosome 8 in a couple in which the male partner is a carrier of a balanced reciprocal translocation (46 XY(11 p14; 14 q21)).

**Table 1.** Incidence of meiotic and mitotic, whole and segmental chromosome abnormalities in trophectoderm biopsies in women above and below 35 years of age.

| Maternal age | No of biopsy samples | Meiotic aneuploidy |  |  |  |  |  |  |  |  | Total abnormalities |
| --- | --- | --- | --- | --- | --- | --- | --- | --- | --- | --- | --- |
|  |  | Paternal |  | Maternal |  | Total | Segmental |  |  | Total |  |
|  |  | Gain | Loss | Gain | Loss |  | Gain | Loss | Complex |  |  |
| ≤34 | 856 | 4 | 17 | 40 | 56 | 117<sup>a</sup> (14%) | 1 | 24 | 6 | 31 (4%) | 148<sup>b</sup> (17%) |
| >35 | 414 | 0 | 6 | 88 | 101 | 195<sup>a</sup> (47%) | 2 | 17 | 4 | 23 (5.5%) | 218<sup>b</sup> (53%) |
| All ages | 1270 | 4 | 23 | 128 | 157 | 312 (25%) | 3 | 41 | 10 | 54 (4%) | 366 (29%) |

<sup>a,b</sup>p=<0.0001

| Maternal age | No of biopsy samples | Mitotic imbalance |  |  |  | Total abnormalities |
| --- | --- | --- | --- | --- | --- | --- |
|  |  | Whole chr | Segmental |  |  |  |
|  |  |  | Single | Complex | Total |  |
| ≤34 | 856 | 64<sup>c</sup> (7%) | 129 | 13 | 142<sup>d</sup> (17%) | 206<sup>e</sup> (24%) |
| >35 | 414 | 42<sup>c</sup> (10%) | 45 | 5 | 50<sup>d</sup> (12%) | 92<sup>e</sup> (22%) |
| All ages | 1270 | 106 (36%) | 174 (58%) | 18 (6%) | 192 (64%) | 298 (23%) |

<sup>c</sup>p=0.1069 <sup>d</sup>p=0.0355 <sup>e</sup>p=0.4684

Segmental aneuploidies with evidence of a meiotic origin were uncommon with only three examples of segmental gain associated with dual maternal haplotypes. In contrast, presumed meiotic segmental losses were more frequent and were predominantly associated with the absence of paternal haplotypes: 58.5% (24/41). For PGT-SR with reciprocal translocations involving terminal segments, combined parental haplotyping and intensity ratio analysis could detect imbalance down to the 5Mb limit of analysis (Fig 2c). However, for confident identification of sporadic segmental imbalances, a consistent change over at least two or three 5Mb regions was required and, to the nearest 5Mb, the affected regions ranged in size from 10 to 130Mb (n=27).

Unlike maternal meiotic aneuploidies, the incidence of presumed mitotic whole chromosome and segmental imbalance, which had parental intensity ratio imbalance equivalent to meiotic trisomies but normal parental haplotype patterns confirming inheritance of single paternal and maternal chromosomes at fertilisation, did not significantly increase with advanced maternal age: 24% 206/856 vs 22% 92/414 (p= 0.4684) (Table 1). Hence, in women aged ≤34 years, the overall incidence of mitotic aneuploidies was greater than those of meiotic origin and conversely in women ≥35 years, meiotic aneuploidies were the predominant type of aneuploidy. Although mitotic whole chromosome imbalance was relatively common, ten samples with 3-7 affected chromosomes accounted for 42 of the 106 of these aneuploidies detected (Fig 2b).

Several samples had multiple karyotype-wide aneuploidies indicating the trophectoderm cells were clonally derived from an abnormal mitotic division and in one case, the combination of multiple monosomies from both parents and two nullisomies is characteristic of a tripolar mitotic division (Ottolini *et al*., 2017). Overall, mitotic whole chromosome imbalance was almost equally split between gain (or loss) of the paternal and maternal chromosomes, 47% 50/106 vs 53% 56/106, respectively. The incidence of mitotic segmental imbalance, either single events and typically terminal imbalances, or complex multiple segmental abnormalities was high and marginally decreased with maternal age (p= 0.0355). Mitotic whole chromosome imbalance was more evenly distributed across all autosomes whereas segmental aneuploidies were more frequent in the larger chromosomes (Figure 3c). The size of the affected regions ranged from 10 to 170Mb (n=55).

**Figure 3.**
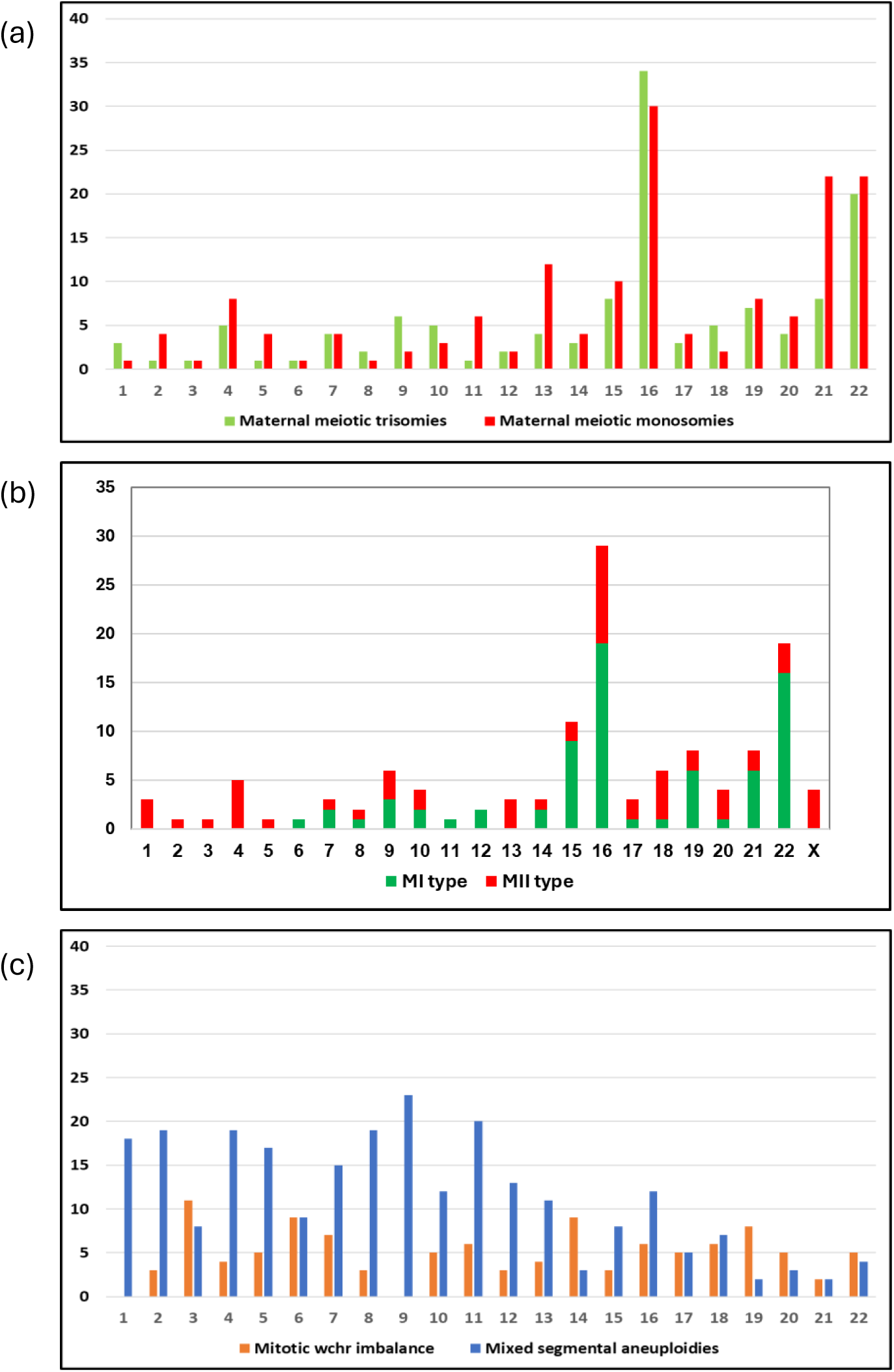
Chromosome distributions of (a) maternal meiotic aneuploidies and (b) origin in the first (MI type) or second meiotic division (MII type) and, (c) mitotic whole chromosome and segmental imbalance (all types).

The number of chromosome abnormalities in individual aneuploid biopsy samples ranged from one to ten, however, 69% (325/471) of samples had only a single abnormality. The aneuploid samples were classified into four groups according to the strength of the evidence for the abnormality and the known severity of clinical outcomes (Table 2; Figure 4): (1) euploid (no abnormalities detected (NAD), (2) mitotic imbalance (whole and segmental), (3) meiotic monosomy (whole and segmental) and (4) meiotic trisomies (whole and segmental). The proportion of aneuploid samples with mitotic imbalance only, decreased marginally from 14% (124/856) to 10% (40/414) (P = 0.0448) in women aged ≤34 and ≥35 years. In contrast, meiotic aneuploidies of all types increased significantly from 17% (142/856) to 40% (165/414) (P <0.0001) in these age groups.

**Figure 4.**
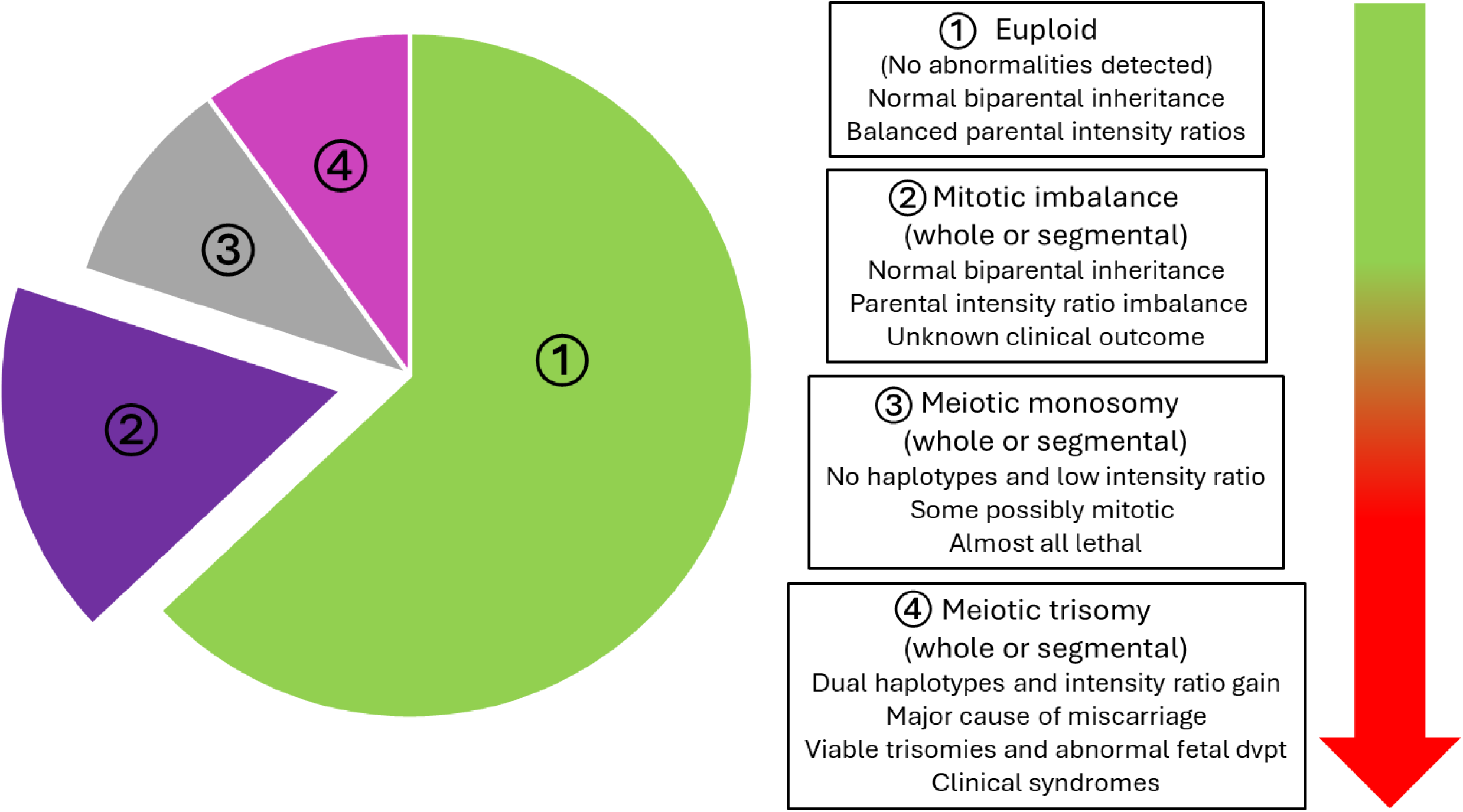
Prioritization scheme for embryo selection based on known adverse clinical outcomes.

**Table 2.** Classification of trophectoderm biopsy samples ranked by severity of known adverse outcomes in women above and below 35 years of age.

| Mat age | No of biopsy samples | Euploid (NAD) | Mitotic imbalance only |  |  | Meiotic aneuploidy (including any additional mitotic imbalances) |  |  |  |  |  |  |  |  | Total abn |
| --- | --- | --- | --- | --- | --- | --- | --- | --- | --- | --- | --- | --- | --- | --- | --- |
|  |  |  | Whole chr | Segmental | Total | Trisomy only | Monosomy only | Mixed | Total | Seg gain | Seg loss | Complex | Total | All meiotic |  |
| ≤34 | 856 | 590<sup>a</sup><br>(69%) | 33 | 91 | 124<sup>b</sup><br>(14%) | 47 | 62 | 4 | 113<sup>c</sup><br>(13%) | 1 | 23 | 5 | 29<br>(2%) | 142<sup>d</sup><br>(17%) | 266<sup>e</sup><br>(31%) |
| ≥35 | 414 | 209<sup>a</sup><br>(50%) | 16 | 24 | 40<sup>b</sup><br>(10%) | 51 | 70 | 23 | 144<sup>c</sup><br>(35%) | 2 | 15 | 4 | 21<br>(4%) | 165<sup>d</sup><br>(40%) | 205<sup>e</sup><br>(50%) |
| All ages | 1270 | 799<br>(63%) | 49 | 115 | 164<br>(13%) | 98 | 132 | 27 | 257<br>(20%) | 3 | 38 | 9 | 50<br>(4%) | 307<br>(24%) | 471<br>(37%) |

<sup>a,c,d,e</sup> P < 0.0001 <sup>b</sup> P = 0.0448

Categorising the cohort of biopsied blastocysts in each cycle by the presence of one or more embryos identified as aneuploid (excluding segmental abnormalities of unknown clinical outcomes), ranked by the likelihood of adverse clinical outcomes shows that the proportion of cycles with all euploid embryos (no abnormalities detected) decreased significantly from 44% (97/218) to 18% (23/124) in women ≤34 and ≥35 years of age, respectively (P <0.0001) (Table 3). Cohorts with only one or more embryos affected by mitotic whole chromosome imbalance of unknown clinical outcome, did not increase with maternal age and were identified in 12% (41/342) of all cycles.

**Table 3.**
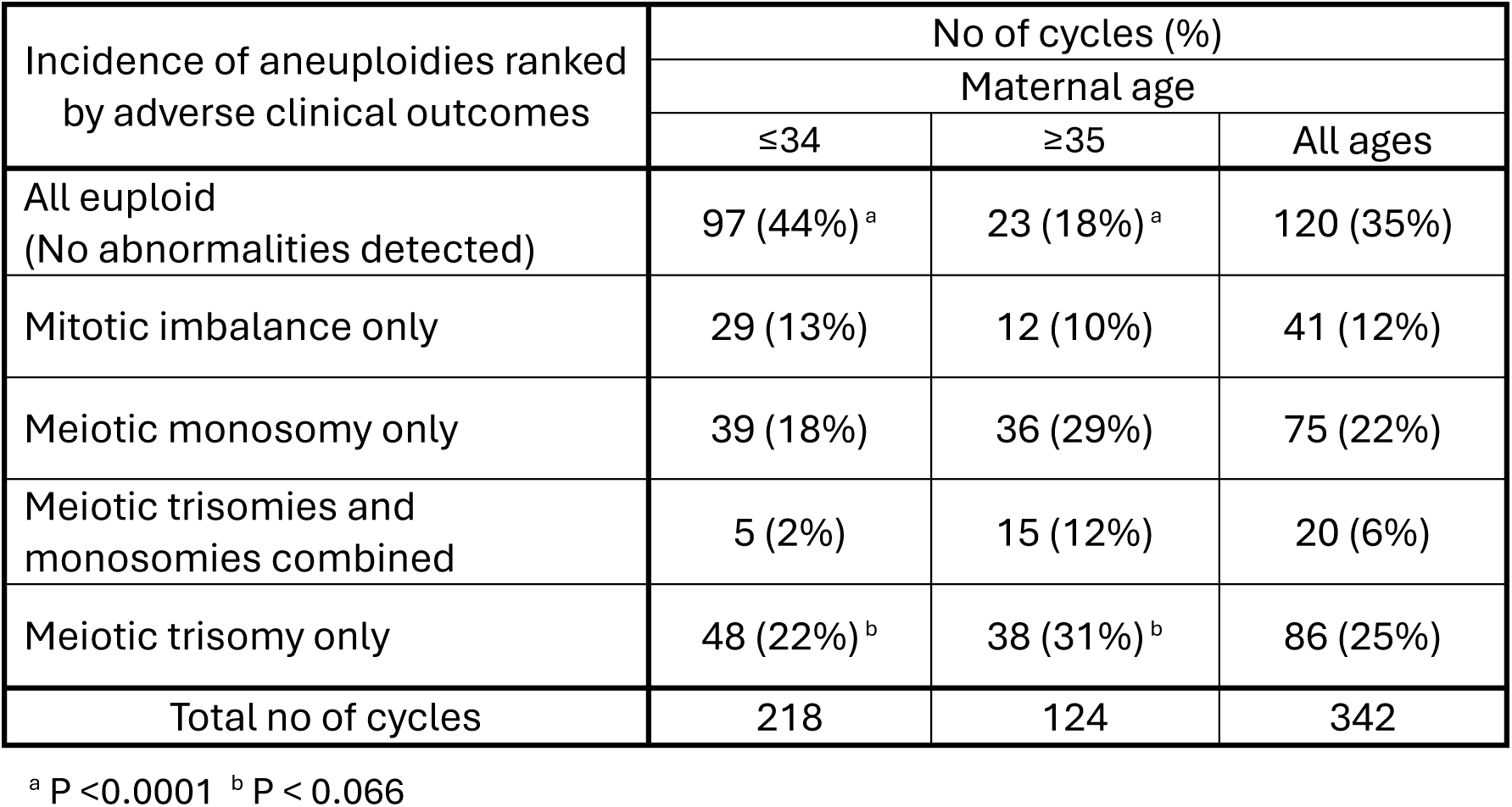
Cycles with one or more blastocysts identified with meiotic or mitotic aneuploidies, ranked by known adverse clinical outcomes.

| Incidence of aneuploidies ranked by adverse clinical outcomes | No of cycles (%) |  |  |
| --- | --- | --- | --- |
|  | Maternal age |  |  |
|  | ≤34 | ≥35 | All ages |
| All euploid<br>(No abnormalities detected) | 97 (44%)<sup>a</sup> | 23 (18%)<sup>a</sup> | 120 (35%) |
| Mitotic imbalance only | 29 (13%) | 12 (10%) | 41 (12%) |
| Meiotic monosomy only | 39 (18%) | 36 (29%) | 75 (22%) |
| Meiotic trisomies and monosomies combined | 5 (2%) | 15 (12%) | 20 (6%) |
| Meiotic trisomy only | 48 (22%)<sup>b</sup> | 38 (31%)<sup>b</sup> | 86 (25%) |
| Total no of cycles | 218 | 124 | 342 |

<sup>a</sup> P < 0.0001 <sup>b</sup> P < 0.066

Cohorts with one or more embryos identified as having meiotic aneuploidies, including trisomies alone, trisomies and monosomies and monosomies alone, all increased with maternal age and collectively increased significantly from 43% (92/218) to 72% (89/124) in these age ranges (P <0.0001).

## Discussion

Genome-wide single nucleotide polymorphism (SNP) genotyping and karyomapping is now well established for preimplantation genetic testing of monogenic disease (PGT-M) (Handyside *et al*., 2010; Natesan *et al*., 2014; Schadwell *et al*., 2022). However, to identify chromosome abnormalities, it has been necessary to examine the karyomap patterns of each chromosome in detail together with log R ratio and B-allele frequency (BAF) plots, which is time consuming and often difficult to interpret, and this has restricted the application of this approach for preimplantation genetic testing for aneuploidy (PGT-A). Using a combination of SNP parental haplotyping and intensity ratio profiling overcomes these limitations in a single low cost assay and has enabled the identification of meiotic and mitotic, whole chromosome and segmental gains and losses together with their parent of origin for comprehensive PGT.

For PGT-A, low-read depth next generation sequencing (NGS) enables high resolution chromosome copy number analysis by mapping sequenced fragments to successive regions or bins across each chromosome (Fiorentino *et al*., 2014). However, with DNA from multiple cell trophectoderm biopsy samples, intermediate copy number changes, between the normal two copies and three copies in trisomies or one copy in monosomies, are common and difficult to interpret. Low level changes may be technical artefacts whereas higher level changes may indicate chromosomal mosaicism within the sample. Furthermore, transfer of embryos with intermediate copy number changes and presumed to represent chromosomal mosaicism has resulted in healthy live births (Greco *et al*., 2015) and with copy number changes below 50% pregnancy rates are similar to euploid embryo transfers (Capalbo *et al*., 2021; Viotti *et al*., 2021, 2023).

Using SNP genotyping and karyomapping for polar body and trophectoderm biopsy analysis in parallel with PGT-A by NGS, we recently demonstrated that although meiotic aneuploidies were, with one exception, always represented in the corresponding trophectoderm biopsy sample copy number profile, some only had intermediate copy number changes whereas some presumed mitotic aneuploidies had full copy number changes (Handyside *et al*., 2021). Thus, applying copy number thresholds is not reliable for distinguishing the important meiotic aneuploidies, which affect the whole embryo, from mitotic aneuploidies arising during preimplantation development and possibly affecting only a variable proportion of the sampled trophectoderm cells. To improve the accuracy of copy number analysis, targeted multiplex amplification or sequence capture methods have been developed to sequence thousands of genome-wide fragments containing SNP loci. This allows both copy number and B-allele frequency analysis. With the targeted approach, a non-selection study confirmed a high positive predictive value for euploid transfers and ongoing pregnancy (Tiegs *et al*., 2021). However, this approach does not distinguish between meiotic and mitotic aneuploidies and the relatively low-resolution limits analysis of segmental aneuploidies.

Combined parental haplotyping and intensity ratio analysis has the disadvantage of requiring both parents to be genotyped. However, SNP microarrays are a low-cost method for consistent genotyping of a large set of defined genome-wide SNPs selected for high heterozygosity ratios and the parental samples only need to be genotyped once. Also, for PGT-A alone, a reference sample from a close relative or sibling embryo is not required to identify regions with dual or missing haplotypes from one parent allowing single samples to be tested, as has been reported previously (Verdyck *et al*., 2022). This allows the full range of meiotic and mitotic chromosome abnormalities to be identified together with the parent of origin. Another approach using NGS for SNP genotyping combined with copy number analysis for PGT-A recently reported similar findings to those presented here (De Witte *et al*., 2024). Furthermore, a method for comprehensive PGT to base pair resolution by whole genome sequencing, haplotyping and haplarythmisis for copy number analysis has been validated (Janssen *et al*., 2024). Thus, there is now an expanding range of laboratory and analytical methods for improved PGT.

For PGT-A, because only a small number of biopsied trophectoderm cells is tested, whatever method is used for aneuploidy detection, the result may not be fully representative of the whole embryo and importantly the inner cell mass of the blastocyst from which the fetus develops (Popovic *et al*., 2020). The exception is meiotic trisomy identified using polymorphic or other markers which detect both homologues from one parent, as demonstrated here by SNP parental haplotyping. As meiotic aneuploidies originate though segregation errors in one or both meiotic divisions during the formation of the gametes and are inherited by the embryo at fertilisation, the whole embryo is affected. Achiasmate bivalent chromosomes have been identified in germ cells in fetal ovaries and particularly affect the smaller chromosomes (Hassold *et al*., 2021). Theoretically, missegregation of achiasmate chromosomes in the second meiotic division could then result in trisomy with two identical chromosomes and therefore a normal single haplotype pattern. Combined parental haplotyping and intensity ratio analysis would identify these events as an imbalance between parental chromosome intensities with no evidence of dual haplotypes resulting in misclassification of the trisomy as mitotic in origin. However, small scale-studies using SNP genotyping of polar bodies and embryo samples with or without NGS-based copy number analysis has not found any evidence that the absence of crossing over causes aneuploidy in mature oocytes (Ottolini *et al*., 2015, 2017; Handyside *et al*., 2021). In this study, 87.5% (28/32) of maternal trisomies for chromosomes 21 and 22 were identified as meiotic and only four were identified as mitotic in origin.

The limitations of testing biopsy samples also apply to identifying meiotic and mitotic monosomies or segmental deletions. As meiotic errors are identified by the complete absence of either haplotype from one parent, it is possible that the chromosome is only absent from the sampled cells and is of mitotic origin. Indeed, there will also be limits to the detection of SNP markers from euploid cells in diploid/aneuploid mosaic samples.

The marginally increased incidence of presumed meiotic monosomies compared with meiotic trisomies, particularly those of paternal origin may indicate that a proportion of presumed meiotic monosomies are mitotic (Table 1). Rebiopsy and analysis of aneuploid blastocysts by comprehensive methods that distinguish different types of aneuploidies should clarify how representative single biopsy samples are of the whole embryo (Cascante *et al*., 2023).

PGT-A has been criticised on the basis that the inaccurate and inconsistent reporting of aneuploidies, often because of NGS-based testing and intermediate copy number abnormalities, can result in the discard of viable embryos, reducing cumulative live birth rates (Paulson, 2023). It is essential therefore to improve methods for accurate detection of aneuploidies, and because of the limitations of testing a single biopsy, to distinguish abnormalities of meiotic and mitotic origin. One strategy to minimise the possibility of eliminating viable embryos, which is made possible by combined SNP parental haplotyping and intensity ratio analysis and other similar methods, is to prioritise embryos for transfer based on the strength of the evidence that an embryo is aneuploid and the known severity of the clinical outcome (Table 3; Figure 4). The classification scheme proposed here is based on increasing evidence of normal live birth outcomes following the transfer of embryos with intermediate, mosaic copy number changes of whole or segmental aneuploidies in trophectoderm biopsies ascertained by NGS-based copy number analysis (Capalbo *et al*., 2021; Viotti *et al*., 2021, 2023; Besser *et al*., 2024). Using combined SNP analysis, it is possible to distinguish between mitotic imbalance with normal biparental inheritance and meiotic aneuploidies with evidence of abnormal inheritance of parental chromosomes and consequently more severe clinical outcomes.

Mitotic whole or segmental imbalances with clear evidence of normal parental haplotype patterns for both the paternal and maternal chromosomes, indicating chromosome mosaicism in an otherwise euploid embryo, were identified in 13% of biopsy samples and this proportion was only marginally lower in women over the age of 35 years (Table 2). Embryos with biopsy samples only affected by these abnormalities, therefore, could be considered for transfer and avoid the discard of potentially viable embryos. The chromosome distribution of mitotic whole chromosome imbalance (Figure 3c) is similar to previous studies using NGS-based copy number analysis, demonstrating that mosaic, intermediate copy number changes, are evenly distributed across all chromosomes (McCoy *et al*., 2023).

With meiotic monosomies, the evidence of meiotic origin is less strong, simply because of the limitations of sampling only a small number of trophectoderm cells. Although there are no informative SNP markers from one parent, associated with a close to zero intensity across the whole of the corresponding chromosome, the loss may not affect all the cells of the embryo, and the clinical outcome is less certain. Indeed, paternal meiotic monosomies are more frequent than paternal trisomies (Table 1). Furthermore, there is recent evidence from single cell analysis that they occur *de novo* late in preimplantation development, possibly confined to the trophectoderm, and consequently only affect a small proportion of cells (Zhai *et al*., 2024). As most meiotic monosomies are lost either before implantation or early in pregnancy, these could therefore be considered for transfer with appropriate genetic counselling, if no euploid embryos were available.

With meiotic trisomies identified by parental haplotyping and intensity ratio analysis, the evidence of meiotic origin is strong, since typically hundreds of informative SNP markers for both homologues from one parent are detected in dual haplotype regions of the chromosome and there is a corresponding intensity imbalance across the whole chromosome (Figure 2a and c). Furthermore, meiotic trisomies, mainly of maternal origin, are known to be a major cause of miscarriage or, in some cases, are viable and can either cause highly abnormal fetal development, or rarely, result in affected live births (Segawa *et al*., 2017; McKinlay Gardner and Amor, 2018; Moorthie *et al*., 2018).

Meiotic segmental gains with evidence of dual haplotypes were rare and all of maternal origin (Table 2). As these may result in viable abnormal pregnancies, they cannot be considered for transfer. In contrast, presumed meiotic segmental losses are more common and are mainly of paternal origin (Kubicek *et al*., 2019; Navratil *et al*., 2020).

Segmental aneuploidies of mitotic origin can arise in the early mitotic divisions following fertilisation (Vanneste *et al*., 2009). Recently, however, single cell analysis of trophectoderm and inner cell mass cells has reported that segmental aneuploidies, a majority of which are losses, are common and often restricted to small numbers of cells, indicating a mitotic origin (Chavli *et al*., 2024). Embryos with biopsy samples only affected by a segmental loss, therefore, could be considered for transfer, depending on the chromosome segment involved. Indeed, normal live births have been reported following transfer of embryos identified with mosaic and non-mosaic segmental aneuploidies (Viotti *et al*., 2021, 2023; Besser *et al*., 2024).

The use of vitrification for cryopreservation of biopsied blastocysts for PGT-A has encouraged the adoption of single embryo transfers, eliminating multiple pregnancies and high implantation and ongoing pregnancy rates per embryo transfer have been reported (Whitney *et al*., 2016; Gorodeckaja *et al*., 2019; Tiegs *et al*., 2021).

Theoretically, however, any type of embryo selection, including PGT-A, cannot increase cumulative pregnancy rates and may reduce rates because of the elimination of viable embryos misidentified as aneuploid (Paulson, 2023). Repeated transfer of single fresh and vitrified-warmed blastocysts should optimise cumulative singleton pregnancy rates from each IVF cycle. However, without PGT-A, the chance of adverse clinical outcomes is also increased. In the cohorts of biopsied blastocysts in each cycle reported here about half had one or more embryos affected by meiotic aneuploidies, divided about equally between samples with meiotic trisomies alone, which are a major cause of miscarriage following IVF (Segawa *et al*., 2017), or monosomies alone, almost all of which are lost before or soon after implantation (Table 3). Thus, in addition to improving ongoing pregnancy rates per transfer, particularly in women of advanced maternal age, PGT-A may have an important role in prioritising euploid blastocysts for cryopreservation and reducing the cumulative rate of miscarriage and other adverse clinical outcomes per cycle following transfer of aneuploid embryos.

## Acknowledgements

The authors would like to thank Dr Michael C. Summers for critically reviewing the paper.

## Authors’ contributions

AHH and JH designed the study. AHH and LN analysed the data and wrote the paper, MN developed the software, DH, JV, JP and JK carried out the lab work and helped with those aspects of the paper and preparation of the figures, JH analysed the data and critically reviewed the paper. All authors had the opportunity to review the final version of the manuscript and make comments and changes before submission.

## Ethical approval

All patients consented to IVF with preimplantation genetic testing by SNP genotyping and karyomap analysis. No new procedures, protocols, or randomization were used in these clinical cases. Consequently, the study does not constitute human subjects research, and accordingly, approval from a Research Ethics Committee was not required.

## Data availability

Patient confidentiality prevents the sharing of the SNP microarray data files. The details of the aneuploidies identified in all samples are provided in Supplementary data.

## Materials and Methods

### Preimplantation genetic testing

Couples having IVF, blastocyst biopsy and preimplantation genetic testing for monogenic disease and/or structural rearrangements by microarray-based SNP genotyping and karyomapping had additional manual reporting of aneuploidy based on examining the karyomapping data for each chromosome in conjunction with logR and B-allele frequency plots as previously published (Natesan *et al*., 2014; Kubicek *et al*., 2019). Briefly, 3-10 trophectoderm cells were biopsied from good quality blastocysts on days 5-7 post insemination. The biopsy samples were lysed and the whole genome amplified (WGA) by multiple displacement amplification (MDA) (REPLI-g Advanced Single Cell Kit, Qiagen, Germany). Parental and reference genomic DNA and the WGA products of the biopsy samples were genotyped by microarray at approximately 300K (HumanKaryomap-12v1.0 BeadChip, Illumina Inc., USA) and 700K SNP loci (Global Screening Array, Illumina Inc., USA) by standard methods and the data analysed using dedicated software (Bluefuse Multi v4.2 and GenomeStudio v2; Illumina Inc, USA).

### Combined genotyping and parental intensity analysis

The SNP data were also analysed using software for karyomapping and to combine parental haplotype and intensity ratio analysis to report meiotic and mitotic, whole chromosome and segmental aneuploidies, using a method which does not require a close relative or sibling embryo to phase the heterozygous SNP loci (Omnia AneuScan®, ExOvo Genomics, UK). For karyomapping, heterozygous and homozygous informative SNP loci in the samples were analysed in successive 1Mb regions across each chromosome and complied into parental haploblocks. Reference and non-reference haploblocks are shown in blue/red and orange/green for the paternal and maternal chromosomes, respectively, purple for regions in which heterozygous informative SNPs for both chromosomes from one parent were present (dual haplotypes) and grey for regions with no informative heterozygous SNPs. For parental intensity ratio analysis, the intensity of paternal and maternal SNP sequence variants were averaged over successive 5Mb regions of each chromosome and plotted as a ratio. The software then combined the data to identify aneuploidies of different types, and these calls were checked manually.

### Categorisation of aneuploidy types

Combined karyomapping and parental intensity ratio analysis enabled meiotic and mitotic, whole chromosome and segmental gains and losses to be identified (For examples, see Figs 2 and 4). Meiotic trisomies were identified by a combination of dual haplotype regions from one parent across regions of the chromosome with a consistent gain in intensity for that parent across the whole chromosome. Whereas mitotic trisomies had a consistent gain in intensity ratio for one parent equivalent to that of a meiotic trisomy without any evidence of regions with dual haplotypes. Meiotic monosomies were identified by a combination of the absence of either haplotype for that parent across the whole chromosome with an intensity ratio approaching zero for that parent. The same criteria were applied to segmental aneuploidies which to be called confidently required a consistent gain or loss from one parent across two or more successive 5Mb segments in the terminal regions of the p or q arms.

### Statistical analysis

The N-1 Chi Squared test was used to compare the percentages of different aneuploidies (MedCalc Ltd; https://www.medcalc.org/calc/comparison_of_proportions.php).

